# Prenatal adversity configures a subpopulation of ventral dentate granule cells for recruitment to drive innate anxiety

**DOI:** 10.1101/2025.02.02.636173

**Authors:** Anika Nabila, Rose Sciortino, Nicole Politowska, Elizabeth Brindley, Peter U. Hámor, Lia Zallar, Judit Gal Toth, Kristen Pleil, Miklos Toth

## Abstract

Adverse prenatal environment is a risk factor for the development of psychiatric disorders. Although studies have implicated epigenetic mechanisms, little is known about how epigenomic changes come about and lead to abnormal behaviors in affected individuals. We sought to identify epigenomic and transcriptomic signatures induced by a proinflammatory gestational environment in the ventral dentate gyrus (vDG), a hippocampal region linked to avoidance of threatening contexts, that persist and promote anxiety-like behavior in mice. Here we show that adversity shifted the methylation of enhancers and promoters with intermediate methylation and altered synapse-related gene expression, resulting in epigenetic and transcriptional heterogeneity in the vDG. Exposure to an anxiogenic environment recruited vDG neurons with the most transcriptional alterations. Differentially expressed synapse-relevant genes in ensemble neurons tended to be differentially methylated. Finally, this ensemble exhibited higher activity in threatening than safe environment suggesting a prenatal adversity-induced epigenetic and neurobiological sequence that leads to anxiety.

## INTRODUCTION

Anxiety disorders are one of the most common mental illnesses with a lifetime incidence of around 30%^1,2^. While avoidance of aversive and threatening environments is an adaptive behavior and promotes survival, exaggerated reactivity to real or perceived threats is maladaptive, manifested as high arousal, vigilance, avoidance, and overt fear-like reactions. Innate anxiety is an intrinsic condition that can develop to generalized anxiety and other forms of anxiety disorders.

Adversity during prenatal and early postnatal life is a significant risk factor for the development of anxiety disorders because of the sensitivity of the fetal and early postnatal development to disruptions^3,4^. Although research has been dominated by childhood adversity and trauma studies, gestational adversity, including maternal stress^5^, maternal infection/inflammation^6^, and maternal diet^7^, represent a major risk factor for anxiety disorders later in life^8^.

In line with human data, gestational adversity results in anxiety-like behaviors in rodents. Maternal immune activation, induced by the viral/bacterial mimetic polyinosinic:polycytidylic acid (PolyI:C) or lipopolysaccharide (LPS), respectively, was reported to cause offspring anxiety-like behaviors such as avoidance of the unprotected open arm of the elevated plus maze (EPM) and the center of the open field (OF), and preference for the relative safety of the closed arm of the EPM and the periphery of the OF^9,10^. Rodent studies that assessed effects of maternal stress during pregnancy reported similar findings^11^. Together, human and rodent studies highlight the significant role of gestational adversity in shaping future anxiety-related behaviors.

An increasing body of evidence supports the notion that adversity induced maladaptive behaviors are driven by epigenetic and associated gene expression changes in the brain^12^. DNA methylation undergoes dynamic changes during sensitive periods of brain development that can be disrupted by environmental insults^13^. Changes in DNA methylation of *NR3C1*, a gene encoding the glucocorticoid receptor involved in stress response, was reported in cord blood cells in association with maternal psychosocial stress and anxiety during pregnancy^14,15^. DNA methylation changes were also found in the promoter of *NR3C1* and *CRH* (corticotropin-releasing hormone also involved in stress response) in the hypothalamus of offspring following maternal stress during pregnancy in mice^16^. An epigenome-wide DNA methylation study identified sex-dependent methylation changes in genes and pathways related to brain development and immune function in the prefrontal cortex^17^. Although these single locus and genome-wide DNA methylation studies revealed the powerful and long-lasting effect of adverse gestational environment on the offspring’s neuronal epigenome, little is known about how these epigenomic changes come about and lead to abnormal behaviors, including anxiety, later in life.

Here we sought to specifically investigate gestational adversity induced changes in DNA methylation and gene expression that can be directly linked to the behavioral manifestation of innate anxiety. We employed a mouse model of prenatal adversity that is timed to the midgestational period. We previously reported that a partial serotonin 1A receptor (5HT1AR) deficit (i.e., heterozygote knockout, KO) in females on the outbred Swiss Webster (SW) background causes peripheral immune abnormality and immune activation^18^. This maternal condition, via the placenta, propagates to the fetal brain and results in increased anxiety-like behavior and heightened stress responsiveness in the genetically wild type (WT) adult offspring^19,20^. Specifically, we reported that offspring anxiety was caused by a midgestational deficit in a chemokine (XCL1) in the maternal circulation, with the concomitant increase in the levels of the proinflammatory molecules TNF, IL-17a, sRAGE, and LCN2 (a proxy for tissue injury) in the fetal placenta^21^ and transmigration of immune cells to the neonatal brain^20^.

Anxiety is a complex assemblage of maladaptive behaviors emerging from highly interconnected prefrontal and limbic regions^22^. A critical contributor is the ventral hippocampus (vHPC), a brain region essential for identifying contexts and attributing their associated threat/aversion levels. There is ample evidence linking high vHPC activity to the manifestation of avoidance of real or perceived threats in an environment. Lesioning the vHPC^23–25^ or inhibiting vHPC projections to the mPFC^26–28^ decreases avoidance of the unprotected open arm of the EPM. Further, increased glutamatergic activity from the basolateral amygdala onto vCA1 pyramidal neurons increases, while reduced activity decreases, EPM avoidance behavior^29^. Finally, direct silencing of the ventral dentate gyrus (vDG) of the vHPC confers resilience to chronic stress-induced anxiety-like behavior (avoidance of the center of the OF), whereas excitation of the vDG promotes susceptibility to anxiety^30^.

The main goal of our study was to identify prenatal adversity induced changes in DNA methylation in vDG granule cells (vDGCs) of the WT offspring of 5HT1AR heterozygote KO mothers that persist into adulthood and, via changes in the neuronal transcriptome and activity, promote the development of innate anxiety-like behavior. We found that genomic regions with an intermediate level of CG methylation and enrichment in annotated transcriptional enhancers and promoters are preferential epigenomic targets of prenatal adversity. Prenatal adversity resulted in a variable impact on the epigenome and transcriptome of vDGCs, with the most affected cells recruited to an EPM induced neuronal ensemble. This ensemble exhibited higher activity in the anxiogenic open than in the safe closed arm, likely contributing to the avoidance of the open arm and to the overall anxiety-like behavior of mice with prior prenatal adversity.

## RESULTS

### Prenatal adversity induced anxiety-like behavior is accompanied by reduced synaptic inhibition and intrinsic excitability in ventral dentate granule cells

We previously reported that the WT offspring of 5HT1AR deficient mothers exhibit anxiety–like behaviors that are moderately sex biased, with males displaying more pronounced EPM anxiety and increased escape directed behavior in forced swim test (interpreted as increased stress responsiveness) than females^19–21^ (**Figure 1A**). Embryo and postnatal crossfostering experiments showed that these behaviors were caused by the pre- and not the postnatal maternal environment^19^. Further, we showed that the offspring behavioral phenotype was due to immune activation in the 5HT1AR deficient female, as well as in the placenta^21^. Therefore, the 5HT1AR deficient maternal model is similar to polyI:C and LPS induced maternal immune activation (MIA) models of prenatal adversity^10,31–38^. Hereafter we refer to the 5HT1AR prenatal adversity model as 5HT1AR-MIA.

**Figure 1:**
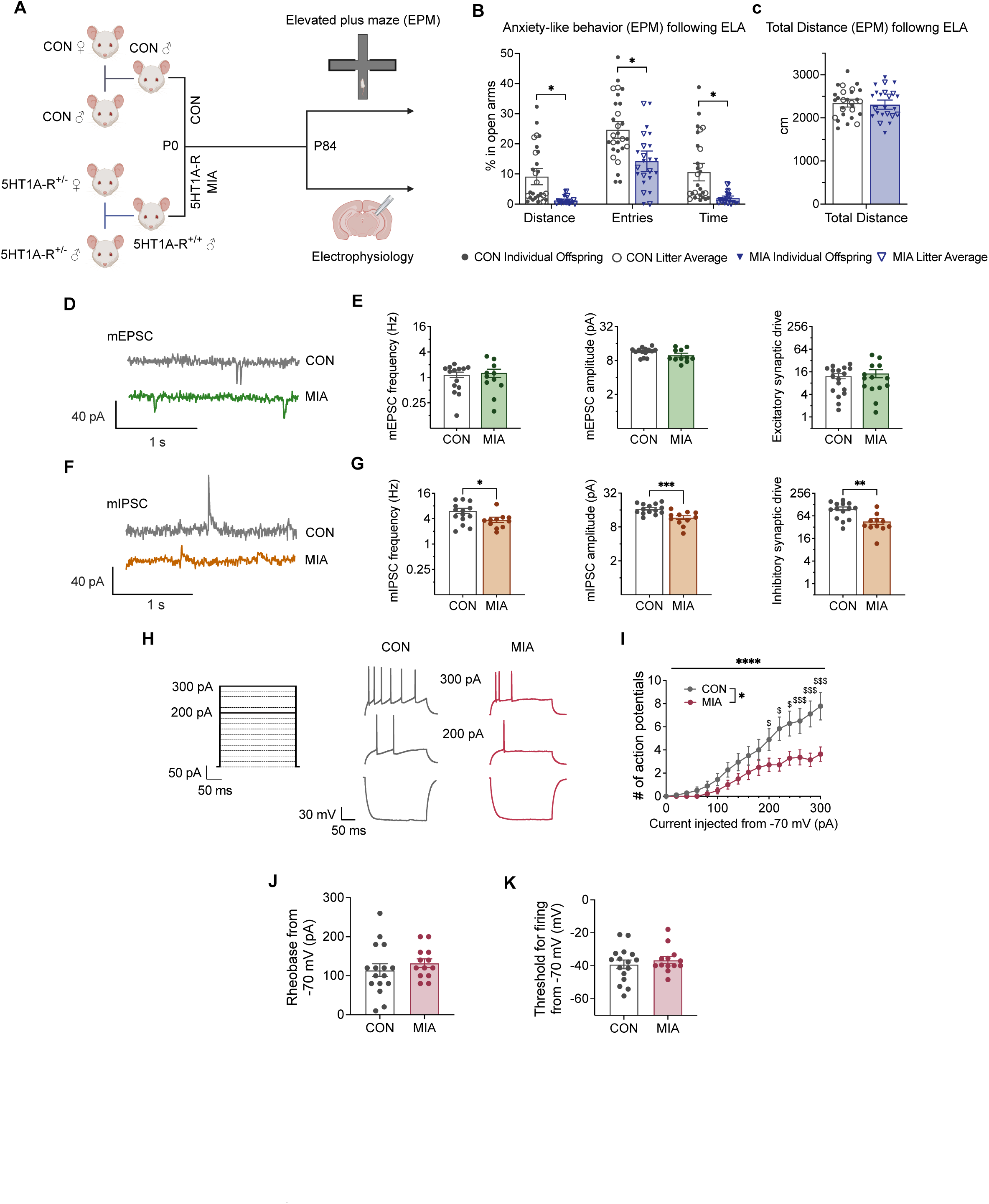
Prenatal adversity results in adult anxiety-like behavior and alters neuronal excitability in vDGCs. **A.** Generation of MIA offspring and experimental design. **B.** Adult MIA offspring (5HT1A-R^+/+^ ♂ offspring of 5HT1A-R^+/-^ parents) covered shorter distance (*p = 0.035), made fewer entries (*p = 0.014) and spent less time in the open arms of the elevated plus maze (EPM) (*p = 0.034) than CON offspring (WT offspring of WT mothers). N = 14 MIA offspring/9 5HT1A-R^+/-^ mothers and18 CON offspring/10 WT mothers. Multiple unpaired t-tests using maternal means/litter averages, corrected with the Holm-Šídák method. Bars are maternal mean ± SEM. **C.** No difference in total distance traveled between MIA and CON offspring. **D-G.** Miniature excitatory and inhibitory postsynaptic currents (mEPSCs and mIPSCs, respectively) in the suprapyramidal blade vDGCs from MIA and CON males. N = 11 cells/5 MIA mice and 14 cells/5 CON mice. (D) A representative trace of mEPSCs. (E) mEPSC frequency, amplitude, and synaptic drive (frequency x amplitude) were not altered in MIA vDGCs (p’s = 0.950, 0.071, and 0.859, respectively). (F). A representative trace of mIPSCs. (G) mIPSC frequency (*p = 0.047) and amplitude (***p = 0.0008) were lower in MIA than CON vDGCs, resulting in decreased inhibitory synaptic drive in MIA than CON vDGCs (**p = 0.001). **H-K.** Intrinsic excitability of vDGCs in MIA and CON mice as measured by current-injected firing in response to increasing 20 pA current steps from a common holding potential of −70 mV (N = 15 cells/4 MIA mice and 18 cells/5 CON mice). (H) Representative traces of action potentials fired in response to 300 and 200 pA steps of current injection stimulus waveform. (I) MIA vDGCs fired fewer action potentials in response to depolarizing current steps than CON vDGCs. 2-way repeated measures ANOVA and post hoc direct comparisons with Holm-Šídák correction for multiple comparisons. *p = 0.0378 for main effect of adversity; ***p < 0.0001 for interaction of current step x adversity; ^$^p < 0.05, ^$$$^p ≤ 0.001 for post-hoc tests. (J, K) There was no difference in rheobase (pA, p= 0.254) and minimum voltage threshold (mV, p= 0.385) needed to elicit firing. Data are shown as mean ± SEM for total cells with individual points. N = 13 MA cells and 16 CON cells. Unpaired t-tests.

We first replicated the anxiety phenotype of the 5HT1AR-MIA male offspring^19–21^. MIA mice spent less time in, made fewer entries into, and traveled a shorter distance in the open arms of the EPM compared to the WT offspring of WT SW mothers, hereafter referred to as control (CON) mice (**Figure 1B**), indicative of increased anxiety-like behavior. There was no change in overall distance traveled (**Figure 1C**), indicating that increased avoidance behavior of the MIA offspring was not due to altered locomotor activity.

Since expression of innate anxiety has been linked to enhanced activity in the vDG^23–30^, we examined the synaptic transmission and intrinsic excitability of vDGCs using whole-cell slice electrophysiology recordings. We found no effect of prenatal adversity on glutamatergic synaptic transmission in vDGCs, as there were no differences in the frequency, amplitude, or overall excitatory synaptic drive (frequency x amplitude) of miniature excitatory postsynaptic currents (mEPSCs) in vDGCs between 5HT1AR-MIA and CON mice (**Figure 1D–E**). In contrast, inhibitory synaptic transmission was decreased by prenatal adversity, as the frequency and amplitude of miniature inhibitory postsynaptic currents (mIPSCs) was lower in MIA than CON vDCGs, leading to reduced inhibitory synaptic drive (**Figure 1F–G**). Together, these results suggest there is an overall disinhibition of vDGCs in MIA mice that may confer an aberrant ability for vDGCs to become activated by excitatory synaptic inputs conveying information about context-specific and behaviorally relevant stimuli associated with anxiety-like behavior^39,40^.

We also examined the effects of prenatal adversity on the intrinsic excitability of vDGCs in adulthood using current clamp recordings. The resting membrane potential (RMP), rheobase, and threshold for firing did not differ between MIA and CON vDGCs (**SFigure 1A, C, D**). However, MIA vDGCs displayed a reduction in the number of action potentials fired across increasing step injections of depolarizing current when measured at both RMP and from a common holding potential of −70 mV (**Figure 1H–K, SFigure 1B**). These results suggest that vDGC excitation may be less sensitive to excitatory synaptic input, perhaps due to activity–dependent plasticity to maintain homeostatic balance in excitability^41^ to compensate for the loss of tonic inhibition by the local interneuron network, indicating a perturbation in ventral hippocampal function.

### Prenatal adversity alters CG methylation at intermediate methylated regions in ventral dentate granule cells

We previously reported prenatal adversity-induced DNA methylation changes in vDGCs collected from adult 5HT1AR-MIA and CON male offspring^20^. Here we repeated this experiment by profiling vDGC methylation from individual adult MIA and CON males to assess both adversity-induced changes and individual variability in DNA methylation. The DGC layer was microdissected from coronal cryostat slices of the vHPC with minimal contamination from other cell types^20,42^ (**Figure 2A**). We isolated vDGC nuclei and used the enhanced version of reduced representative bisulfite sequencing (eRRBS)^43,44^ that expanded the number of CGs profiled for DNA methylation to ∼3 million at 10x coverage. The four biological replicates (from individual animals) in the MIA and CON groups showed high concordance in methylation, and hierarchical clustering separated the two groups indicating that prenatal adversity resulted in changes in the vDGC methylome that persisted to adulthood and were consistent across individual mice (**SFigure 2A**).

**Figure 2:**
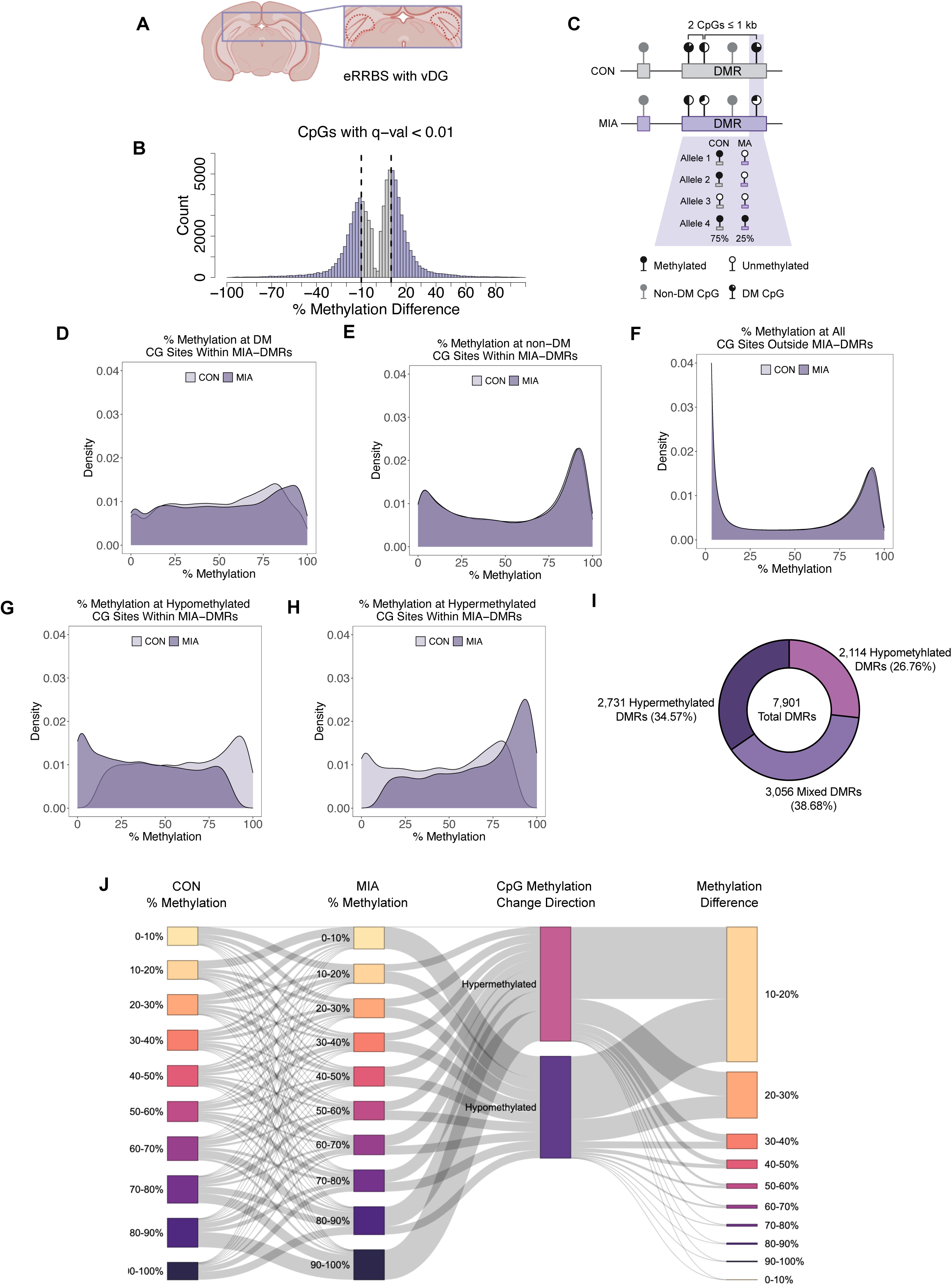
Prenatal adversity preferentially alters methylation at intermediately methylated CGs in vDGCs. **A.** The granule cell layer of the ventral HPC that contains the cell bodies of vDGCs was microdissected for DNA methylation profiling. Dashed line traces the dissected granule cell layer. **B**. Distribution and frequency of methylation differences between MIA and CON at CGs that passed the FDR q< 0.01 threshold. Dashed lines at 10% methylation difference separate DM (purple) and non-DM (gray) CGs. **C**. Two or more DM CGs within 1 kb of each other are clustered to DMRs. Open and closed lollipops indicate unmethylated and methylated CGs, respectively. Intermediate methylation is caused by methylation heterogeneity across cells, represented schematically by one CG in four MIA and CON cells. **D-E**. Methylation distribution of DM (D) and non-DM (E) CGs within DMRs. **F.** Methylation distribution of CGs outside of DMRs. **G-H**. Methylation distribution of hypo- and hypermethylated DM CGs in MIA and CON DGCs. **I.** Proportions of hypomethylated, hypermethylated and mixed categories of DMRs. **J**. Sankey diagram summarizing trends of methylation shifts between MIA and CON DM CGs.

Of the 2,904,392 CG sites mapped in all replicates at >10x coverage, 2.75% of CGs underwent methylation change (at FDR q <0.01) by prenatal adversity, with a slight bias toward hypermethylation vs. hypomethylation (**Figure 2B**). For differential methylation (DM) between MIA and CON DGCs, we set the minimum to 10% (**Figure 2B, SFigure 2B**) because this threshold has been reported to balance sensitivity and specificity in DM detection across RRBS analyses methods^45^. Further, we clustered 2 or more CGs within 1 kb of each other as coordinated changes in CG methylation in a region may have larger regulatory potential^46^ (**Figure 2C**). Overall, 23,676 (0.8%) of CGs had a methylation difference of >10% (at q<0.01) and clustered regionally (<1kb between CGs), specifying 7,901 DM regions (DMRs, 2.67 CGs on average).

Notably, DM CGs within DMRs, both in CON and MIA DGCs displayed mostly intermediate methylation (IM, 10% – 90% methylation) (**Figure 2D**). IM at a given IM CG, or a cluster of coordinately regulated CGs, corresponds to the percentage of the methylated epiallele in a cell population (**Figure 2C**). Since this value also represents the percent of cells carrying the methylated epiallele, DGCs exhibit DNA methylation heterogeneity, especially given the combinatorial effect of the methylated and unmethylated epialleles at thousands of DMRs. In contrast, CGs outside of DMRs showed a genome typical bimodal methylation distribution as they were either uniformly unmethylated or methylated in DGCs (**Figure 2F**). Within DMRs, DM CGs were interspersed with “non-DM” CGs at a 1:2.2 ratio. Non-DM CGs showed partial IM distribution indicating that many of them have methylation bistability like DM CGs in the vDGC population (**Figure 2E**). Of note, the wide range of IM of DM CGs in CON mice indicated that methylation bistability was not due to the mixture of the DGC and non-DGC cell populations such as the small and steady population of progenitors or young (<3-week-old) neurons present in adult DGs^47^.

Analyzing hyper- and hypomethylated DMR DM CGs separately showed that their native, generally IM state (in CON DGCs, **Figure 2G, H**, light purple curves) typically remained in the IM range after prenatal adversity (**Figure 2G, H**, dark purple curves). Further, neighboring DM CGs tended to show a similar directionality of change as ∼60% of DMRs had either all hypo or all hypermethylated DM CGs (26.76% and 34.57%, respectively) (**Figure 2I, SFigure 3**). DMRs with mixed hypo- and hypermethylated DM CGs had twice the size of hypo/hypermethylated DMRs (300.1, 160.0, and 158.3 bp, respectively) but had a similar average number of DM CGs indicating larger distances between CGs that may disrupt their coordinated response to prenatal adversity (**SFigure 3**). Indeed, methylation of CGs in partially methylated regions was reported to be coordinated only when in close proximity (<15 bp)^48^, which was typically seen in DMRs.

The average adversity induced methylation change was ∼22% for both hyper- and hypomethylated DM CGs but varied from the 10% threshold to 30%, with few demonstrating larger changes (**Figures 2B, J**). Given the coordinated methylation changes at hypo/hypermethylated DMRs, MIA decreased and increased the fraction of the methylated DMR epiallele at hypo- and hypermethylated DMRs, respectively. Consequently, the proportion of vDGCs that carry the methylated epiallele of an individual DMR was also altered. For example, a DM CG or their cluster at a DMR with a 50% baseline methylation (in CON) and 20% adversity induced hypomethylation means that the fraction of DGCs harboring the methylated epiallele of the particular DM CG/DMR was reduced from 50% to 30% by prenatal adversity.

IM sites gradually attain methylation from an unmethylated state in progenitors during postnatal DGC maturation^13^. The proinflammatory environment in the fetal and early postnatal brain of MIA offspring may influence this dynamic by setting a lower or higher IM set point than the baseline methylation state in CON vDGC, thus generating the hypo- and hypermethylated DM CGs/DMRs. Interestingly, lower than 60-70% baseline methylation was associated with a higher IM set point and thus producing hypermethylated DM CGs/DMRs in MIA DGCs, while higher baseline methylation with lower IM set point produced hypomethylated DMRs (**Figure 2J**). Overall, these data indicate that prenatal adversity switches the methylation state of individual DMRs from methylated to unmethylated or vice versa in some cells. It is not clear if all or most DMRs undergo epiallelic switching in the same subpopulation of DGCs or different but overlapping subpopulations of DGCs. Nevertheless, the combinatorial effect of adversity induced methylation switching at thousands of DMRs is expected to be variable within the population of DGCs, with some cells affected more than others.

### Regions hypomethylated by prenatal adversity are enriched in cis-regulatory chromatin states

Approximately two-thirds of MIA DMRs were located within the gene body (**Figure 3A**, inner circle), a significant enrichment, due to the overrepresentation of exons, compared to the distribution of randomized control regions (**Figure 3A**, outer circle; Exons Odds Ratio=1.28, p=7.32E-22, Introns OR=0.94, p=1.38E-02). Although around one third of DMRs were intergenic, they were relatively depleted in this genomic feature (OR=0.84, 1.20E-12), suggesting that MIA DMRs tend to be located in protein-coding regions of the mouse genome rather than in intergenic regions.

**Figure 3.**
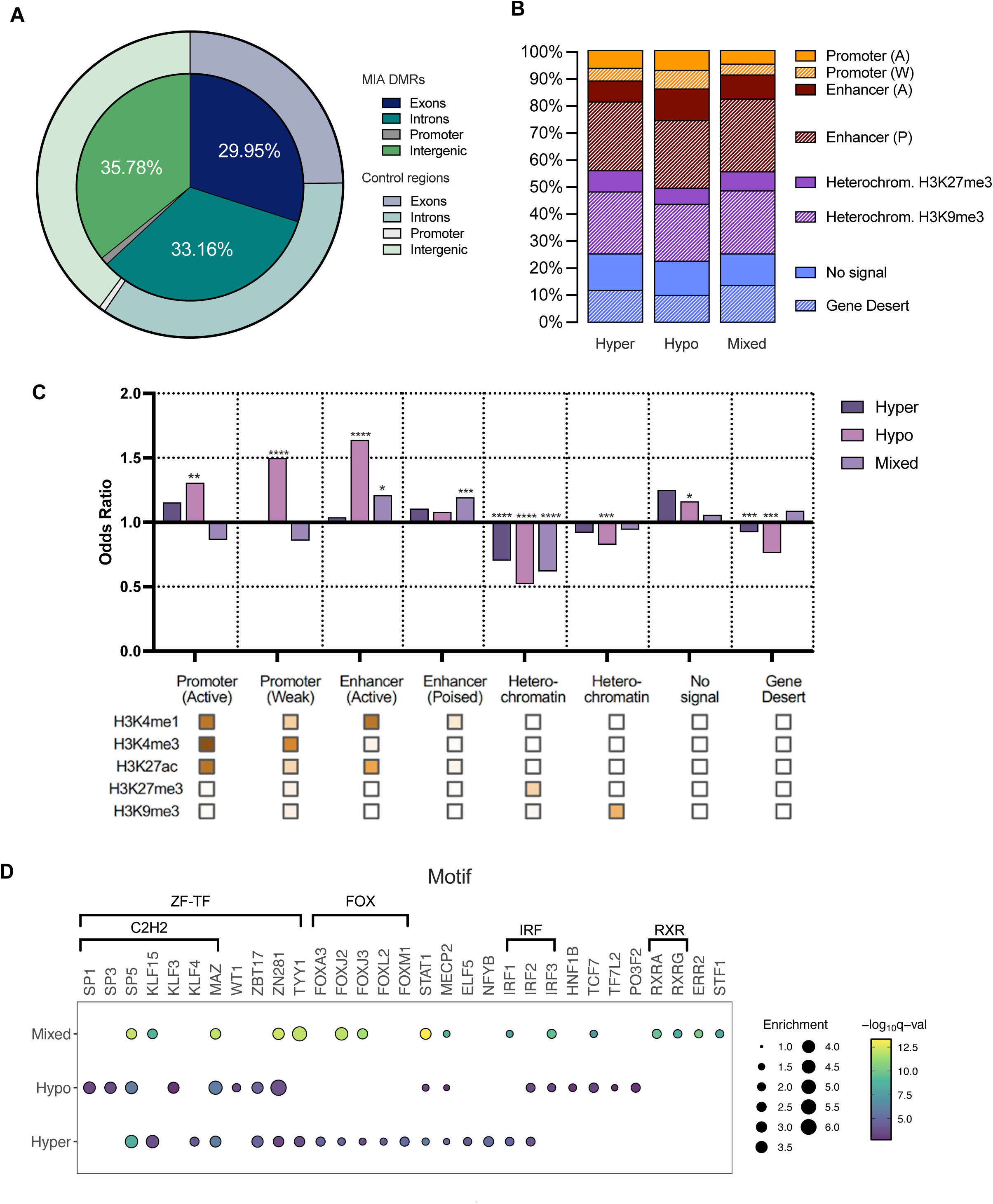
5HT1AR-MIA DMRs are enriched in putative enhancer and depleted in heterochromatin states. **A**. Distribution of MIA DMRs across gene features as compared to randomized control regions. **B**. Distribution of MIA DMRs in ChromHMM^54^ chromatin states, as determined in mouse DGCs^55^. **C**. Enrichment/depletion of MIA DMRs in HMM chromatin states. Enrichments were calculated using Fisher’s exact test and multiple testing correction using false discovery rate (FDR < 0.05), comparing to background block regions clustered from all tested CGs. ****FDR<0,0001, ***FDR<0,001, **FDR<0,01, *FDR<0,05. **D.** Enrichment (E-value < 0.05) of transcription factor motifs, ordered by class, in hypomethylated, hypermethylated, and mixed DMRs identified by SEA from the MEME Suite tools. The top 15 TFs in each category are selected and corresponding TFs with q<0.05 in all categories and their enrichment and q-values are displayed in.

IM is often associated with enhancers^49–53^, therefore we assigned ChromHMM^54^ chromatin states, as previously determined in mouse DGCs^55^, to MIA-DMRs. ChromHMM categorizes chromatin into distinct states (enhancer, promoter, heterochromatin etc.) based on the combinatorial presence and absence of histone modifications. Of the 7,901 MIA DMRs, ∼35% were in an enhancer-specific chromatin state based on the distinct combinations of the histone marks H3K4me1, H3K4me3, and H3K27ac in CON DGCs (**Figure 3B**). Because the basal methylation state of DMRs (in CON DGCs) may influence their chromatin association and the chromatin state the directionality of adversity induced methylation change, we analyzed hypomethylated, hypermethylated, and mixed DMRs separately. Hypomethylated and mixed DMRs, due to their slightly higher numbers, were enriched in enhancer chromatin states; in particular, hypomethylated DMRs were overrepresented in active chromatin (**Figure 3C**). Similarly, hypomethylated but not hypermethylated and mixed DMRs were enriched in active and weak promoters. Although a substantial fraction of DMRs was associated with facultative and, in particular, constitutive heterochromatin signatures (HK27me3 and H3K9me3, respectively), DMRs were depleted in heterochromatin. This is consistent with the wide distribution of heterochromatin across the genome and the reduced accessibility, thus epigenetic malleability of these regions to adversity induced perturbations (**Figure 3B, C**). Overall, a substantial fraction of DMRs (45-50%) was associated with promoters and enhancers that reached significant enrichment with hypomethylated DMRs and, to a lesser extent, with mixed DMRs indicating the potential gene regulatory role of MIA-DMRs.

### Distinct sets of transcription factor binding sites in hypo- and hypermethylated DMRs

The different baseline methylation of hypo- and hypermethylated DMRs (in CON animals) and their different directionality of change by MIA suggested distinct sequence contexts and binding sites for sequence-specific transcription factors (TFs)^56^. We used MEME Suite^57^ to discover putative TF binding motifs within DMRs. We found that although hypo- and hypermethylated DMRs shared a number of putative TF binding motifs, they had an overall dissimilar motif-pattern (**Figure 3D**). First, although they shared C2-H2 zinc finger (ZF) TF binding motifs^58^, such as SP5 and MAZ, binding sites for additional SP family members were present in hypomethylated DMRs. Further, the two types of DMRs contained sites for different KLF family members. SP/KLF/MAZ bind to similar but distinct GC rich targets and function as repressors or activators depending on their interacting co-regulators^59^. MAZ has been reported to interact with CTCF to control cohesin positioning and genome organization and may act as an insulator protein^60^. Binding site for WT1, a ZF TF, was present in hypomethylated DMRs only. Binding of KLF4^61^, SP1^62,63^, and WT1^64^ are DNA methylation sensitive, thus adversity induced switching in a given cell from the unmethylated to the methylated epiallele or vice versa can alter TF binding and possibly DMR promoter and enhancer activity. Second, the cognate binding motif for YY1 (TYY1), still another ZF TF and which is linked to stress sensitivity^65^, was uniquely enriched in hypermethylated DMRs. Similarly, binding sites for four forkhead box (FOX) TFs that can act as pioneer factors to initiate gene transcription in condensed chromatin during large scale chromatin remodeling^66^ were exclusively in hypermethylated DMRs. Finally, mixed DMRs contained many of the same TFs but also unique binding sites. This included, most prominently, unique sites for two members of the RXR nuclear receptor family (RXRA and RXRG) which are part of the RAR/RXR heterodimers that transduce retinoid signaling during development and in neuronal stress and neuroinflammatory processes in adult brain.

### Prenatal adversity configures the transcriptome of a subpopulation of ventral dentate granule cells for activation in an anxiogenic environment

Adversity induced epigenomic changes are expected to lead to transcriptomic changes that may explain the increased anxiety phenotype of MIA mice in the EPM test. Because of ensemble coding in the DG^67^, gene expression of cells that are recruited and activated in an anxiogenic environment is the most relevant to the anxiety-like behavior of MIA mice. Therefore, we transcriptionally profiled DGCs that were activated in the EPM, separately from the rest of the DGCs of CON and MIA males. We used the expression of the immediate early gene (IEG) product FOS, a proxy for neuronal activation^68^, to differentiate activated from not activated cells by fluorescent activated nuclei sorting. Adult MIA and CON mice were exposed to the EPM for 10 min and the vHPC was isolated one hour later for nuclei isolation. Then, FOS+ and FOS- vDGC nuclei (also labeled by the DGC marker PROX1^69^) were sorted for single nucleus (sn) RNA-Seq (**Figure 4A**). A small and comparable fraction of nuclei were FOS+ in CON (0.85%) and MIA (0.98%) mice (average from two experiments), consistent with sparse coding and the tight regulation of ensemble size by inhibitory interneurons in the DG^70–72^. A total of 16,456 MIA FOS-, 10,576 CON FOS-, 3,797 MIA FOS+, and 1,575 CON FOS+ nuclei were obtained from two replicates of 6 MIA and 6 CON mice each and used, after filtering, for clustering and expression analyses.

**Figure 4.**
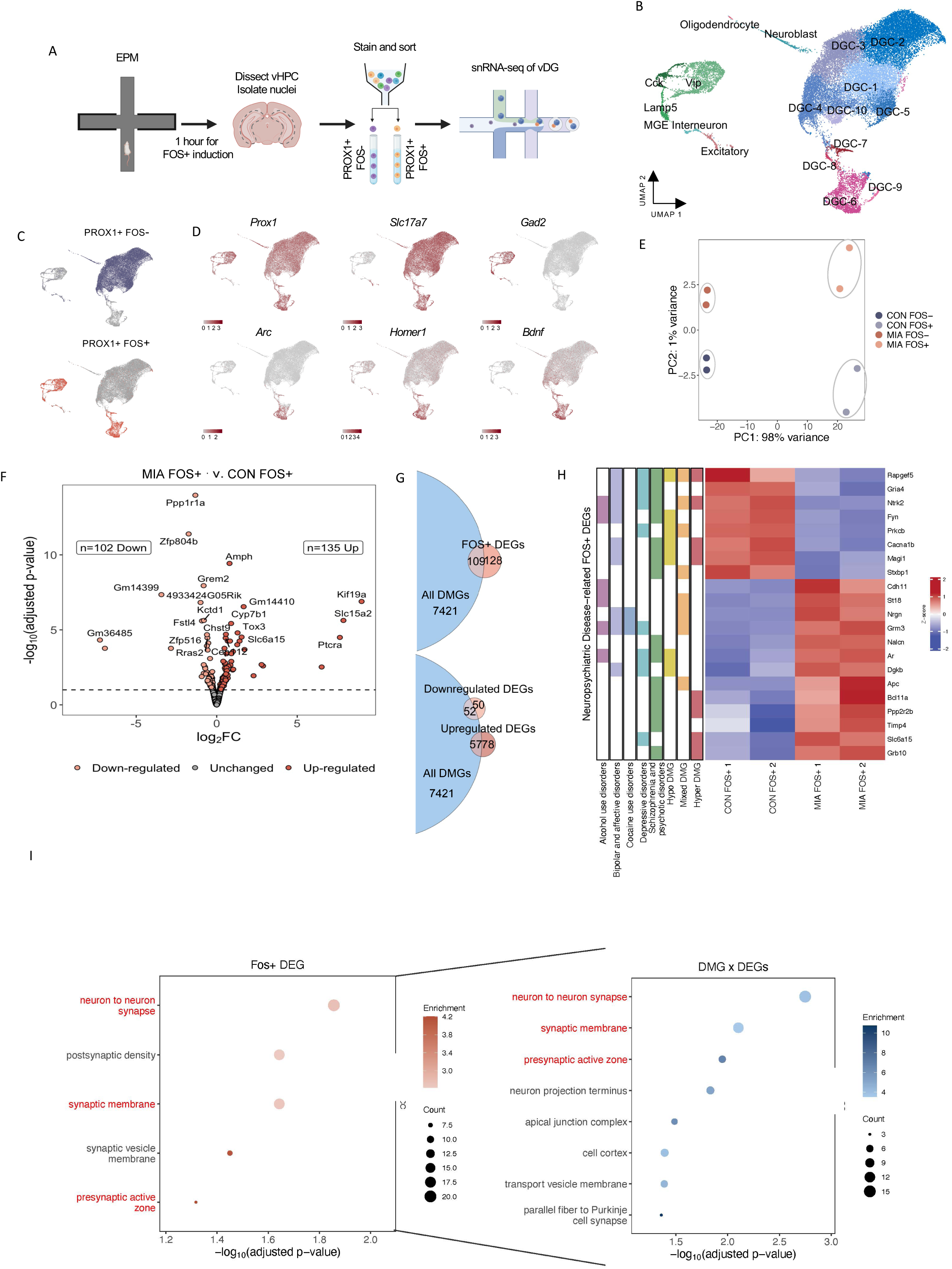
MIA is associated with more transcriptional changes in FOS+ than FOS- vDGC nuclei in adult mice. **A.** Schematic of snRNA-seq experiment. Mice were placed on the EPM after which activated (FOS+) and non-activated (FOS-) vDGC nuclei were isolated for RNA sequencing. N = 2 replicates, each pooled from 6 animals per group. **B.** UMAP of clustered FOS+ and FOS- nuclei. **C.** UMAP of c lusters colored by sample type: FOS- (purple, top) or FOS+ (orange, bottom). **D.** Expression of the marker genes for DGCs (*Prox1*), excitatory neurons (*Slc17a7*), inhibitory neurons (*Gad2*) and activated neurons (*Arc*, *Homer1*, *Bdnf*) in different clusters. **E.** Principal component analysis of pseudobulk CON FOS-, MIA FOS-, CON FOS+ and MIA FOS+ samples. Two replicate samples represent each group (6 mice per sample). **F.** Volcano plot of differentially expressed genes (DEGs, Wald’s test, p-adjusted < 0.1, dashed line) in FOS+ MIA vs. CON vDGC nuclei. **G.** GO “cell compartment” overrepresentation analysis of FOS+ DEGs (left) and FOS+ DEGs x DMGs (right). Red lettering indicates shared GO categories. All significant (p-adjusted < 0.05), non-redundant categories are visualized. **H.** Overlap between DMR genes (assigned to DMRs by GREAT^61^) and DEGs. **I**. Heatmap depicting z-scored expression of FOS+ DEGs that are also neuropsychiatric risk genes. Genes are color coded based on DMR type and neuropsychiatric disease classification.

Nuclei from all conditions were combined and assigned to 17 cell clusters that were separated based on cell-type marker expression and neuronal activation (**Figure 4B, SFigure 4, STable 1**). Clusters DGC-1 to DGC-5 and DGC-10 nuclei formed a large assortment of closely related clusters that contained nuclei primarily from FOS- samples, while nuclei in the smaller DGC-6 to DGC-9 clusters were derived from FOS+ samples (**Figure 4B, C**). In addition, smaller clusters of neuroblasts and oligodendrocytes were mostly derived from FOS- samples while non-DGC excitatory cells and various interneurons (Cck, Lamp5, Vip and MGE interneurons) were derived largely from FOS+ cells. Although PROX1 is considered a marker for DGCs, its expression in interneurons and oligodendrocytes (**Figure 4D**, *Prox1*) is consistent with previous reports^73^. All DGC clusters expressed the excitatory neuronal marker *Slc17a7,* and all interneurons expressed the inhibitory neuronal marker *Gad2* as expected (**Figure 4D**, *Slc17a7, Gad2*). Additionally, clusters that were composed of FOS+ DGCs expressed the IEGs *Arc*, *Homer1* and *Bdnf* (**Figure 4D**). As previously documented, FOS+ interneurons did not strongly express IEGs^74^. Finally, all clusters expressed a similar number of genes and transcripts and were composed of similar proportions of nuclei in CON and MIA samples, including the DGC clusters (**SFigure 4**), indicating that adversity does not lead to overt changes in cell identity and in global transcription in the adult DG.

Next, we combined individual DGC clusters into FOS- and FOS+ DGC pseudobulk samples to assess transcriptomic differences between MIA and CON DGCs. Although most variation in the dataset was related to whether MIA and CON cells were activated or not in EPM (**STables 2** and **3**), likely representing preexisting differences^75^, principal component analysis (PCA) of pseudobulk samples also showed separation between CON and MIA mice for both FOS- and FOS+ DGCs, indicating underlying differences in gene expression due to prenatal adversity (**Figure 4E**, PC1 and PC2, respectively). Surprisingly, both replicates of FOS+ samples displayed a more pronounced separation (e.g., in PC2) between MIA and CON compared to FOS- samples, indicating that EPM- activated vDGCs have greater differences in adversity induced gene expression. Indeed, we observed 135 upregulated and 102 downregulated genes (i.e., FOS+ DEGs) in MIA compared to CON FOS+ DGC nuclei (**Fig. 4F, STable 4**). In contrats, only 31 upregulated and 29 downregulated genes (FOS- DEGs) were identified (**SFigure 5A, STable 5**) in MIA compared to CON FOS- DGC nuclei, 34 of which were also FOS+ DEGs (**SFigure 5B**). Differential expression in non-DGC nuclei that included various interneurons, neuroblasts, and oligodendrocytes (**Figure 4B**) could not be analyzed because of individually small populations. Overall, these data show that DGCs with the most adversity induced transcriptomic changes are preferentially activated in future stressful anxiogenic environments.

FOS+ DEGs were enriched in synapse related genes (gene ontology (GO) cellular components) relevant to both the presynaptic active zone (at mossy terminals) and postsynaptic density (at the entorhinal input) (**Figure 4G** left panel**, SFigure 6**) implicating altered synaptic functions at multiple levels in MIA DGCs. FOS- DEGs, on the other hand, showed enrichment in only one category, neuron projection terminus (p-adjusted = 0.037) (**SFigure 5C**).

Although close to 50% of the 237 FOS+ DEGs (both up- and downregulated) were associated with DM genes (i.e., 7,530 DMGs, assigned to DMRs by GREAT^61^), most methylation changes and DMRs were not associated with expression changes (**Figure 4H**). Nonetheless, the smaller group of 109 FOS+ DEG x DMGs were enriched in several of the same synapse related GO cell compartment categories as the FOS+ DEGs (synapse, synaptic membrane, and presynaptic zone). Furthermore, DEGs within the synapse related GO categories were more likely to be associated with DMRs than genes not enriched in GO categories (Yates corrected X^2^(1,N=237)=4.3705, p=036567). FOS- DEGs x DMGs were represented by only 34 genes, 18 of which were also in the group of the 109 FOS+ DEGs x DMGs, highlighting again the more robust transcriptional response in FOS+ than in FOS- cells by adversity.

### Adversity induced differentially expressed genes are associated with neuropsychiatric disease risk genes

Because anxiety is a common symptom across psychiatric diseases and anxiety disorders are often comorbid with other mental health conditions, such as depression, post-traumatic stress disorder (PTSD), obsessive-compulsive disorder (OCD), and panic disorder, we investigated whether FOS+ DEGs were associated with known neuropsychiatric disease risk genes. Using the PsyGeNET database (version 2.0, 1,549 genes and 117 psychiatric disease concepts)^76^, we found that 21 FOS+ DEGs (9%) were previously implicated in neuropsychiatric disease (**Figure 4I**). Further, 11 of these 21 (∼52%) genes were also DMGs, showing concurrent perturbations in methylation and transcription by prenatal adversity in this set of genes. Some of these genes were common to multiple diseases (e.g., *Ntrk2* in alcohol use disorders, bipolar and affective disorders, depressive disorders and schizophrenia and psychotic disorders) consistent with the transdiagnostic nature of many psychiatric disease risk genes^77^. Interestingly, a majority (14 genes, ∼67%) of DEGs were represented specifically in schizophrenia and psychotic disorders, which is consistent with maternal gestational infection/immune activation being a risk factor for schizophrenia^78,79^ and with maternal polyI:C induced MIA causing a schizophrenia-like phenotype in mouse offspring^80,81^. Overall, the data reveals a relationship between DNA methylation and transcriptional regulation of synaptic genes and neuropsychiatric disease, specifically in FOS+ vDGCs following prenatal adversity.

### Ventral dentate granule cells of mice with prenatal adversity exhibit increased calcium activity in threatening relative to safe environment

Epigenomic and transcriptomic profiling indicated a variable impact of prenatal adversity across vDGCs with the most adversity-affected neurons allocated to the FOS+ activated ensemble during an EPM trial. To assess the in vivo activity of this relatively small but behaviorally most relevant population of DGCs under anxiogenic challenge, we recorded the population level Ca^2+^ signal^82^, a proxy for neuronal activity, in the vDG during EPM performance. This was achieved using fiber photometry monitoring of the virally-expressed fluorescent Ca^2+^ sensor GCaMP6s (**Figure 5A**). The average frequency of normalized vDG GCaMP activity in MIA mice was greater when mice were in the open arms of the EPM, the “anxiogenic” compartment of the maze, than when they were in the relative safety of the center or closed arms, while frequency in CON animals was comparable across the three compartments of the EPM (**Fig. 5B-C**). These data are in line with the increased avoidance of MIA mice and the positive correlation between vDG activity and avoidance behavior^23–30^. There was no difference in the average amplitude of normalized GCaMP activity between MIA and CON animals and across the compartments of the EPM (**Fig. 5B, D**).

**Figure 5.**
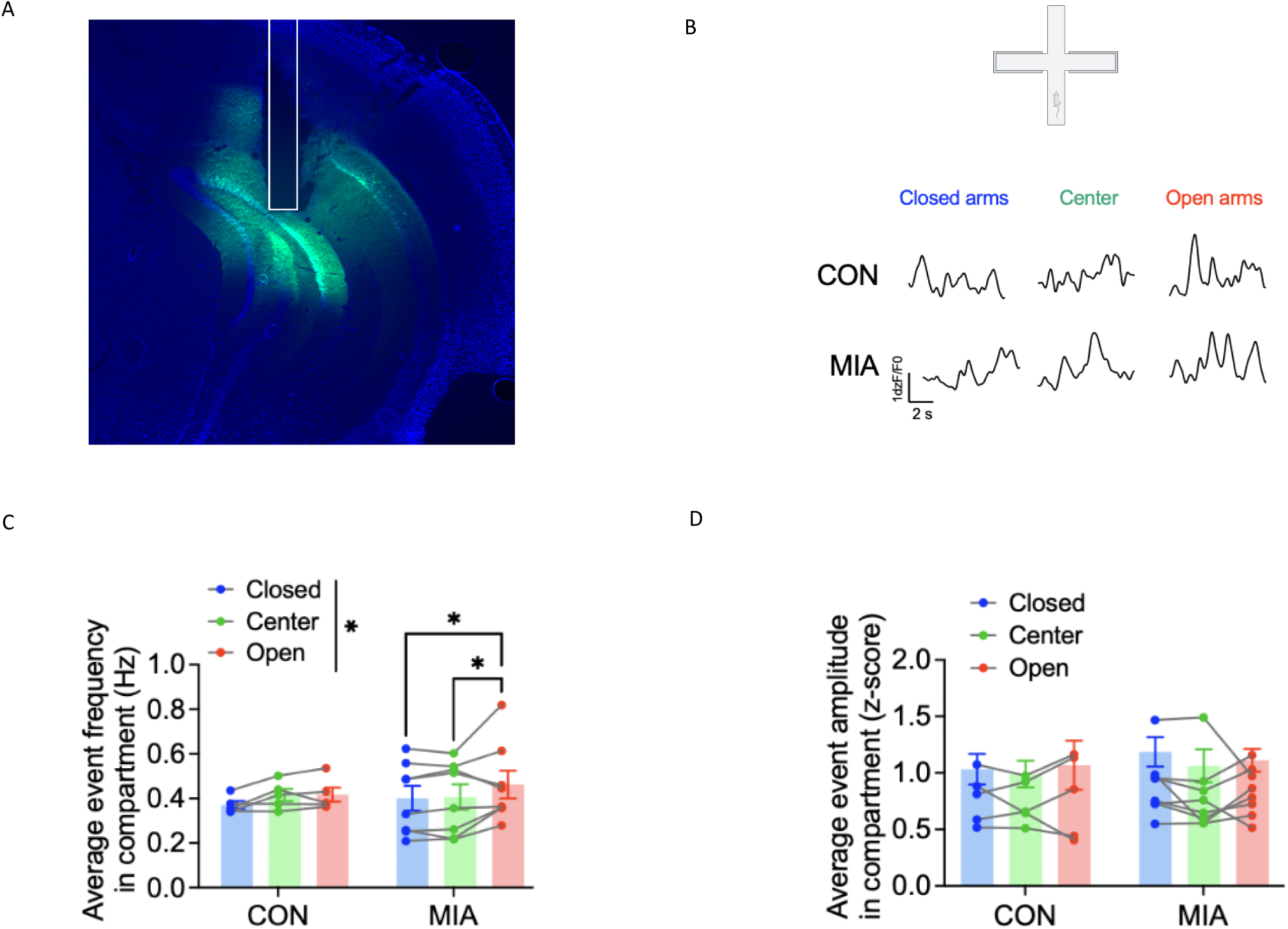
GCaMP fiber photometry in vDG of MIA and CON mice during EPM. **A**. Representative image of GCaMP expression and optical fiber placement in the vDG of a CON mouse. **B**. Representative GCaMP traces from CON (top) and MIA (bottom) in the closed arms, center, and open arms of the EPM. **C**. Greater average frequency of GCaMP transient events in the open arms of the EPM than the closed arms or center, particularly in MIA mice (N=8 MIA and 5 CON mice). 2xRM-ANOVA (main effect of EPM compartment: F(2,22) = 4.12, p = *0.030, no effect of group (p=0.772) or interaction (p=0.346). Post hoc direct comparisons with Holm-Sidak corrections between compartments within group (MIA: open vs. closed *0.0449 and open vs. center *0.0487). **D**. The average amplitude of GCaMP transient events did not differ between compartments of the EPM or between groups. 2xRM-ANOVA: no effect of compartment (p=0.607) or group (p=0.627) or interaction (p=0.808).

## DISCUSSION

The main finding of our study is that prenatal adversity is associated with DNA methylation changes at enhancers and promoters, and alters gene expression in synapse relevant functions in adult vDGCs and that vDGCs with the most pronounced transcriptomic changes are preferentially recruited to a FOS+ ensemble in the stressful environment of the EPM. Further, ensemble activity was higher in a threatening relative to safe environment, consistent with the increased avoidance behavior of MIA mice. Our data suggest for the first time an epigenetic and neurobiological sequence that connects prenatal adversity to later life anxiety-like behavior. This mechanism may provide a plausible explanation for the lifelong increase in stress response and avoidance behavior of individuals with innate anxiety.

### Adverse gestational environment preferentially targets intermediately methylated CGs

Prenatal adversity, represented by the proinflammatory intrauterine environment of 5HT1AR heterozygote mothers, resulted in methylation changes in thousands of small genomic regions with an intermediate level of CG methylation. The level of IM at individual DM CGs and DMRs is presumably set by the underlying sequence and chromatin environment. Specific DNA sequence features were shown to play a role in establishing methylation at partially methylated domains (PMDs)^48^, although PMDs are much larger genomic areas (average of 153 kb) than adversity DMRs (150-200 bp) and are found in only a subset of cell types (cultured fibroblasts and cancer cells). This suggests that sequence alone is not sufficient to establish IM. Since active and bound TFs can dynamically modulate methylation at cis-regulatory elements^50,83,84^, chromatin and associated TFs^56^ are likely involved in maintaining the IM state at adversity DMRs.

### Adversity induces methylation switching at DMRs

Coexistence of the methylated and unmethylated epialleles in the same population of cells suggests relatively unobstructed epiallelic switching that could be sensitive to environmental influences like the proinflammatory environment in the developing MIA brain^20^. We reported that not only the maternal peripheral immune system and the placenta but also the brain of newborn pups exhibit signs of inflammation in the MIA offspring^20^ (see also Introduction).

The 10- 30% (mean 22%) difference at most DMRs in bulk methylation between MIA and CON DGCs indicates epiallelic switching at a given DM CG/DMR in a similar proportion of vDGCs by the adverse experience. Since the fetal hippocampus at the time of prenatal adversity (during the midgestational period^21^) consists of proliferating progenitors^85,86^, and because IM in young DGCs is established by preferential methylation after the first week of life^13^, adversity induced epiallelic shifts likely occur during DGC maturation in the MIA brain. Further, DMRs with higher and lower than the 60-70% baseline methylation in CON DGCs become hypo- and hypermethylated in MIA brains, as a result of lowered and increased IM setpoint, respectively.

### Epigenetic malleability of cis regulatory elements by adverse gestational environment

IM regions are often in an enhancer specific chromatin state^49–53^ and ∼35% DMRs were associated with enhancer-specific chromatin in CON DGCs. An additional 10-15% of DMRs were in promoter specific chromatin states. Overall, these data suggest that up to 50% of the epigenomic targets of prenatal adversity in the 5HT1AR MIA model are IM regions with enhancer or promoter specific chromatin associations and thus can have a regulatory activity.

### The most adversity affected cells are allocated to a FOS+ ensemble in an anxiogenic environment

Consistent with sparse coding in the DG^71,72^, only a small fraction of vDGCs was activated and expressed the activity marker FOS during exploration in the EPM. Remarkably, FOS+ DGCs had four times more adversity induced DEGs than FOS- DGCs, indicating that cells with the most transcriptomic changes are preferentially activated during an EPM task. It is unlikely that the higher number of adversity-induced DEGs in the FOS+ DGC population was due to the EPM exposure one hour prior nuclei isolation because none of the DEGs (between MIA and CON) were IEGs. Rather, adversity induced differential expression between MIA and CON DGCs predated the EPM trial and likely contributed to the allocation of cells with the most adversity induced gene expression changes to the FOS+ ensemble in MIA mice. Finally, FOS+ DEG were exclusively enriched in synapse relevant GO categories and DEGs assigned to these categories were more likely than unassigned DEGs to be associated with DMRs. This suggests a role of DNA methylation in the adversity induced differential expression of some synapse relevant genes, but most DMR genes were not differentially expressed in agreement with the notion that DNA methylation is just one of the epigenetic mechanisms regulating gene expression.

### Prenatal adversity-induced epigenomic and transcriptomic maladaptation in the ventral dentate gyrus governs future anxiety behavior

Why are the most adversity-affected DGCs allocated to the EPM activated ensemble? Since relative excitability has been shown to be a major determinant of neuronal allocation to a coding ensemble in the DG^88^, as well as in the CA1 region and amygdala^89,90^, cells that are most affected by adversity in the epigenetically and transcriptionally heterogenous population of MIA vDGCs may have increased excitability. However, expression of FOS and other IEGs was comparable in MIA and CON FOS+ cells (e.g., were not DEGs) suggesting no apparent difference in neuronal activity between the MIA and CON FOS+ DGC populations. Another factor in determining neuronal allocation is functional connectivity. It has been shown that preexisting connectivity promotes allocation of CA1 neurons to a coding ensemble^90^.

Similarly, CA3 neurons with a similar birth date are functionally connected as neurons born earlier and later during embryonic development are recruited to different ensembles during fear conditioning in adults ^91^. Because adversity resulted in expression changes in pre and postsynaptic genes, it is reasonable to hypothesize that the most transcriptionally altered MIA DGCs have increased connectivity and are preferentially activated in the EPM. Further, given the correlation between vDG activity and innate anxiety, the enhanced in vivo activity of the EPM-activated ensemble in the anxiogenic open arm, relative to the safe closed arm, in MIA mice is consistent with their increased avoidance behavior. Taken together, we propose that prenatal adversity exerts a variable impact on the epigenome and synaptic transcriptome of vDGCs. The most affected vDGCs, which display changes primarily in synaptic genes, are preferentially recruited in an anxiogenic environment, and their heightened activity contributes to avoidance and anxiety-like behaviors.

## Acknowledgements

We would like to thank the support of the Epigenomics Core and the Genomics Core of Weill Cornell Medicine. We are grateful for grant support 5R01MH117004 and 5R01NS106056 to MT and R01AA027645 to KP.

## Author contributions

AN, RS, NP, PH, EB, LZ, and JGT conducted the experiments

AN, EB, NP, PH, KP, and MT designed the experiments

AN, KP, and MT wrote the paper

## Supplemental information

Figures S1–6

## Data Availability

GEO accession number for sequencing data.

DNA methylation by RRBS

GSE284416: https://www.ncbi.nlm.nih.gov/geo/query/acc.cgi?acc=GSE284416

Token: orsvcwygljwhbah

SnRNA-Seq

GSE284463: https://www.ncbi.nlm.nih.gov/geo/query/acc.cgi?acc=GSE284463

Token: wzijwosetxspbir

