## Supplemental Figures for "Prenatal adversity configures a subpopulation of ventral dentate granule cells for recruitment to drive innate anxiety"

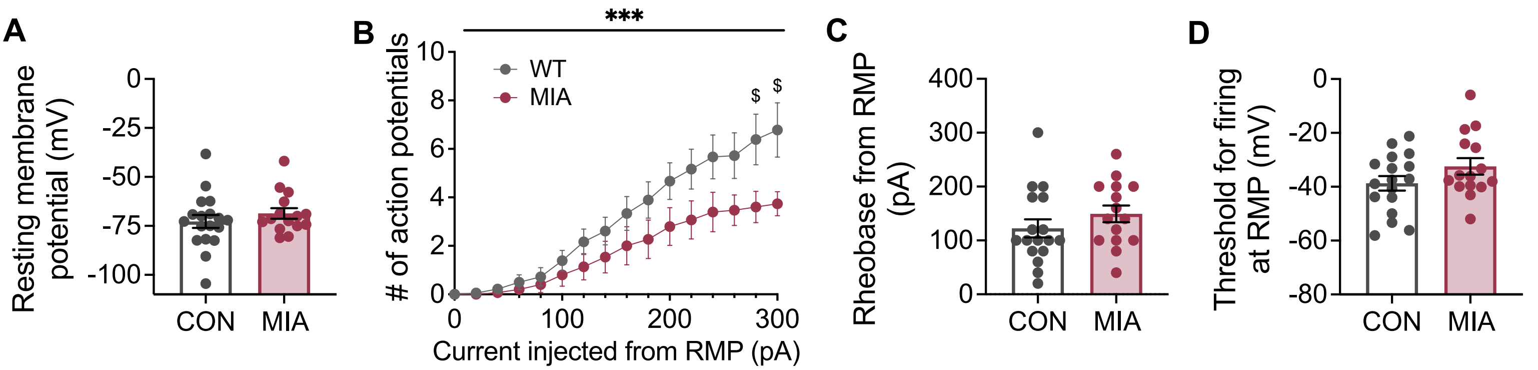


**SFigure 1. Intrinsic excitability of vDGC at resting membrane potential (RMP, related to Fig. 1). A.** The RMP of CON (n = 18 cells) and 5HT1AR-MIA vDGCs (n = 15 cells) did not differ (p>0.05). **B.** MIA vDGCs fired fewer action potentials in response to increasing 20 pA steps of current injection form RMP. 2-way RM-ANOVA with post hoc Holm-Šídák's multiple comparisons test: ***p = 0.0003 for group x current step interaction, $^{\$}p < 0.05$ as indicated for in post-hoc t-tests between group within step. **C, D**. There was no difference in the rheobase (pA) or action potential threshold (mV) between CON and MIA vDGCs (ps > 0.05). Data are shown as mean ± SEM with individual data points where possible.

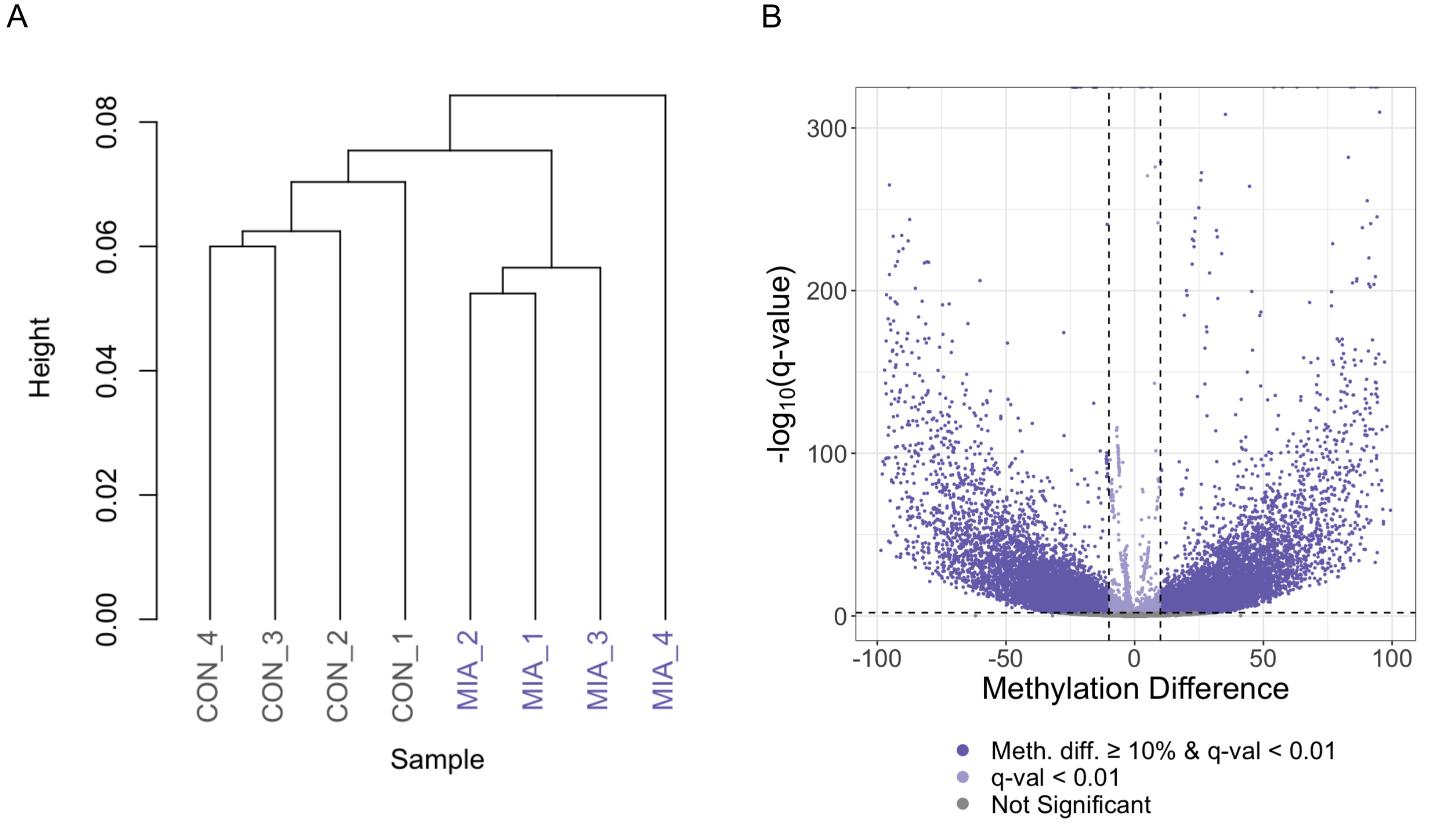


**SFigure 2**. **DNA methylation in vDGCs from individual 5HT1AR-MIA and CON males**. **A**. Hierarchical clustering (1-Pearson's correlation as distance, Ward's method) of CON and MIA replicates (n = 4). **B**. Volcano plot visualizing the methylation difference in all CG sites tested for differential methylation (logistic regression with SLIM method) between MIA and CON. Dots indicate CG sites that passed the threshold for differential methylation (≥ 10% and FDR q-value < 0.01).

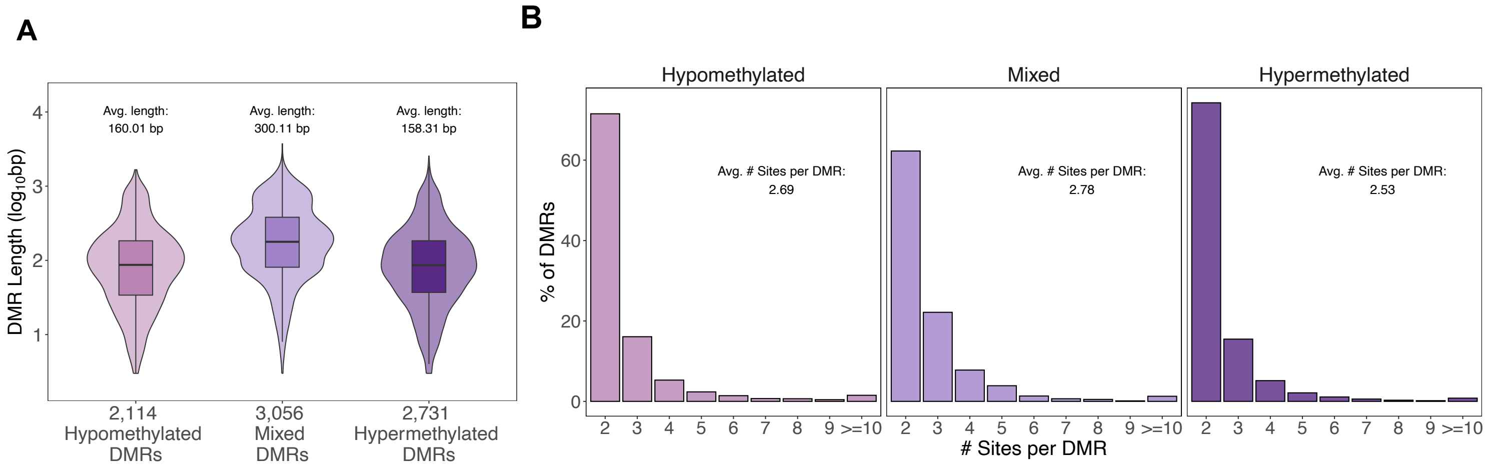


**Sfigure. 3. Characterization of 5HT1AR-MIA DMRs**. **A**. Boxplots showing distribution of DMR length at each DMR category. **B**. Distribution of the number of sites per DMR.

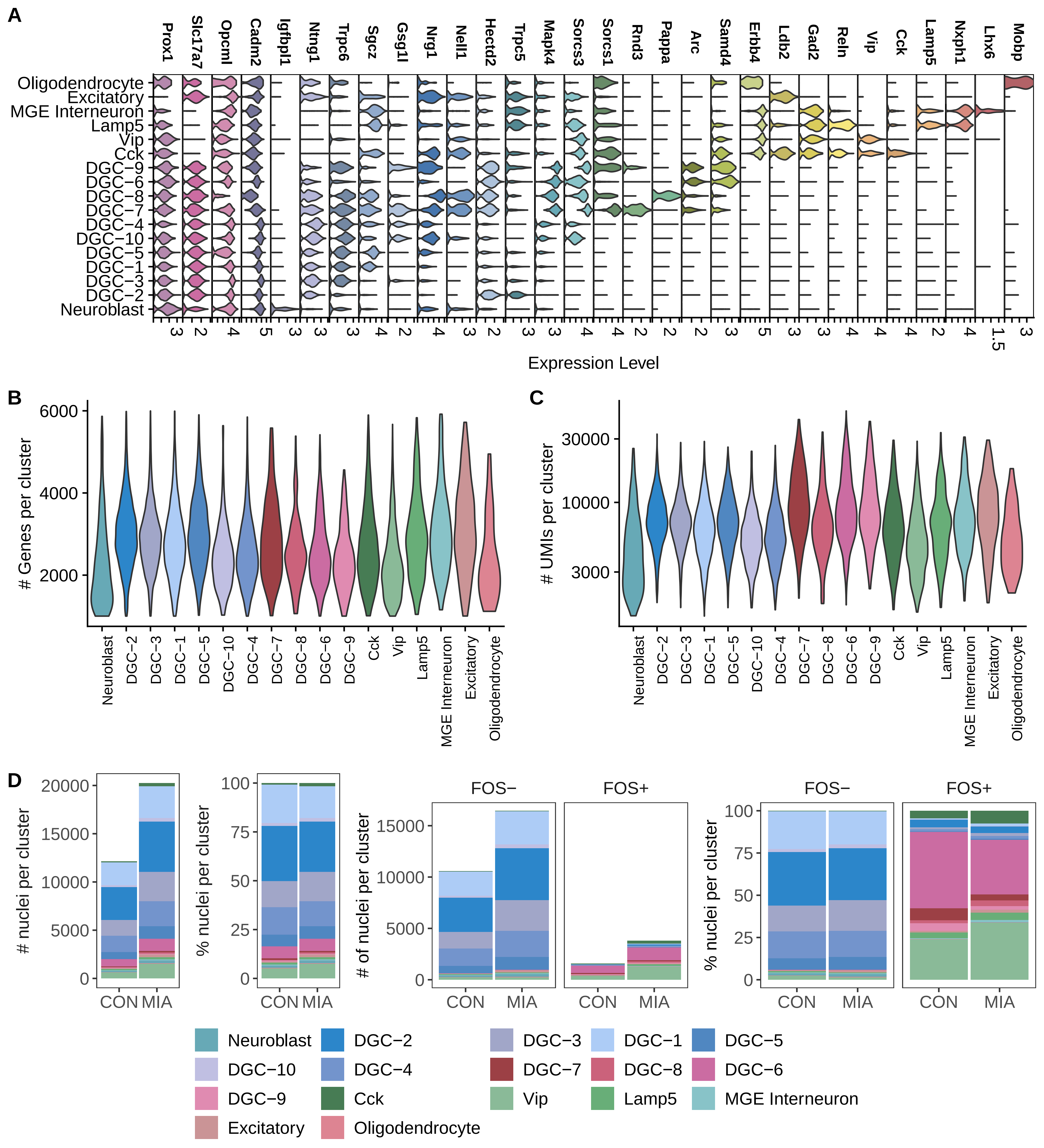
A
Prox1
Slc17a7
Opcml
Cadm2
Igfbpl1
Ntng1
Trpc6
Sgcz
Gsg1l
Nrg1
Nell1
Hectd2
Trpc5
Mapk4
Sorcs3
Sorcs1
Rnd3
Pappa
Arc
Samd4
Erbb4
Ldb2
Gad2
Reln
Vip
Cck
Lamp5
Nxph1
Lhx6
Mobp
Oligodendrocyte
Excitatory
MGE Interneuron
Lamp5
Vip
Cck
DGC−9
DGC−6
DGC−8
DGC−7
DGC−4
DGC−10
DGC−5
DGC−1
DGC−3
DGC−2
Neuroblast
Expression Level
B
### Genes per cluster
6000
4000
2000
C
### UMIs per cluster
30000
10000
3000
Neuroblast
DGC−2
DGC−3
DGC−1
DGC−5
DGC−10
DGC−4
DGC−7
DGC−8
DGC−6
DGC−9
Cck
Vip
Lamp5
MGE Interneuron
Excitatory
Oligodendrocyte
D
### nuclei per cluster
% nuclei per cluster
### of nuclei per cluster
FOS−
FOS+
CON
MIA
Neuroblast
DGC−2
DGC−3
DGC−1
DGC−5
DGC−10
DGC−4
DGC−7
DGC−8
DGC−6
DGC−9
Cck
Vip
Lamp5
MGE Interneuron
Excitatory
Oligodendrocyte

**SFigure 4. Quality control of snRNA-seq data. A**. Marker gene expression in each cluster. **B, C**. Violin plots showing distribution of the number of genes per cluster (B) and the number of UMIs per cluster (C). **D**. Number and proportion of nuclei per cluster of all nuclei together or broken down by FOS-/FOS+ samples.

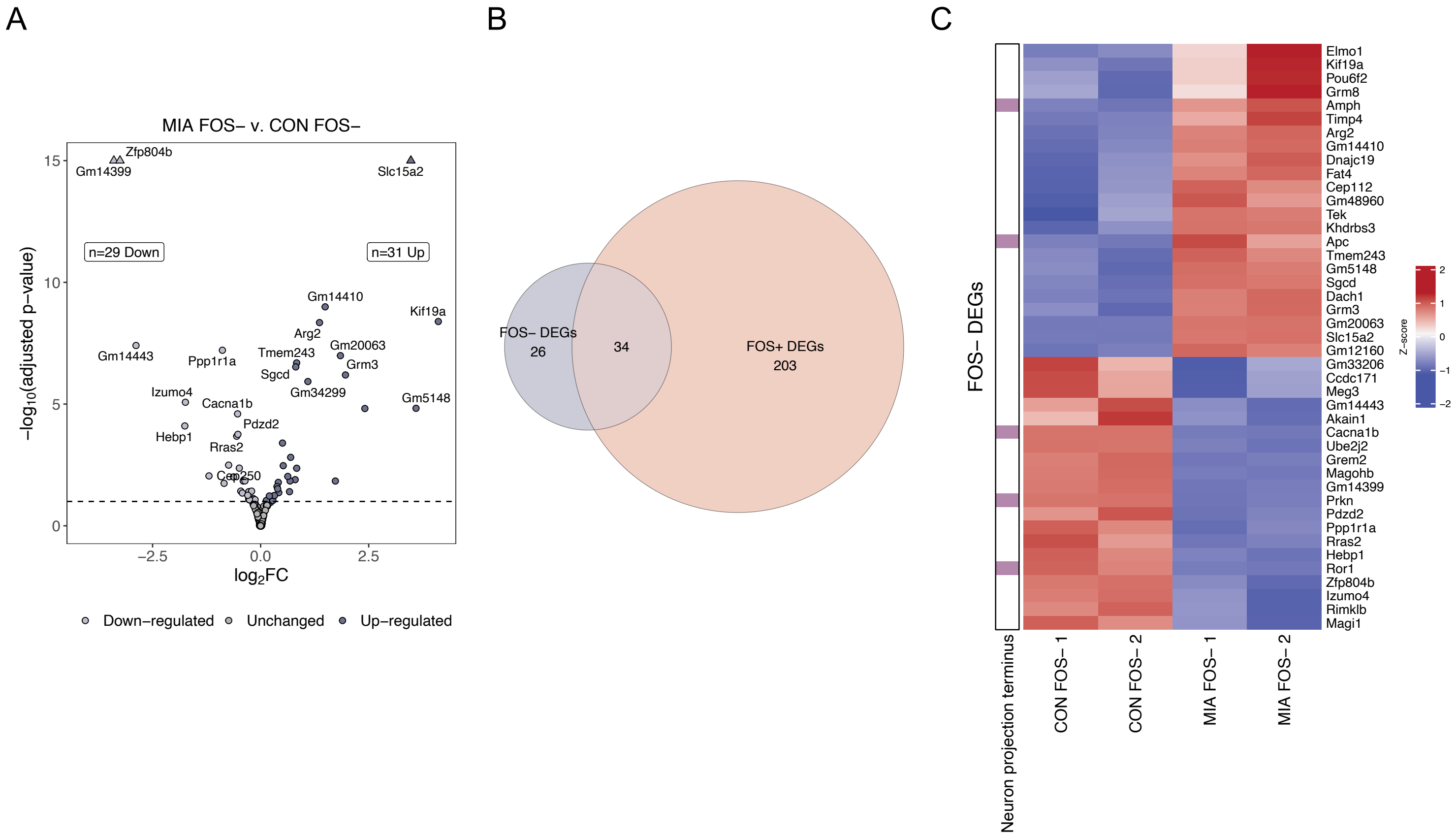


S**Figure 5. FOS- MIA vDGCs exhibit modest changes in transcription. A**. Volcano plot of differentially expressed genes (DEGs, Wald's test, p-adjusted < 0.1, dashed line) in FOS- MIA vs. CON vDGCs. Genes with -log10 p-adjusted > 15 were set to 15 and denoted with a triangle for visualization. **B**. Overlap of FOS- DEGs with FOS+ DEGs. **C**. FOS- DEGs are enriched in only one GO category: neuron projection terminus (p-adjusted = 0.037).

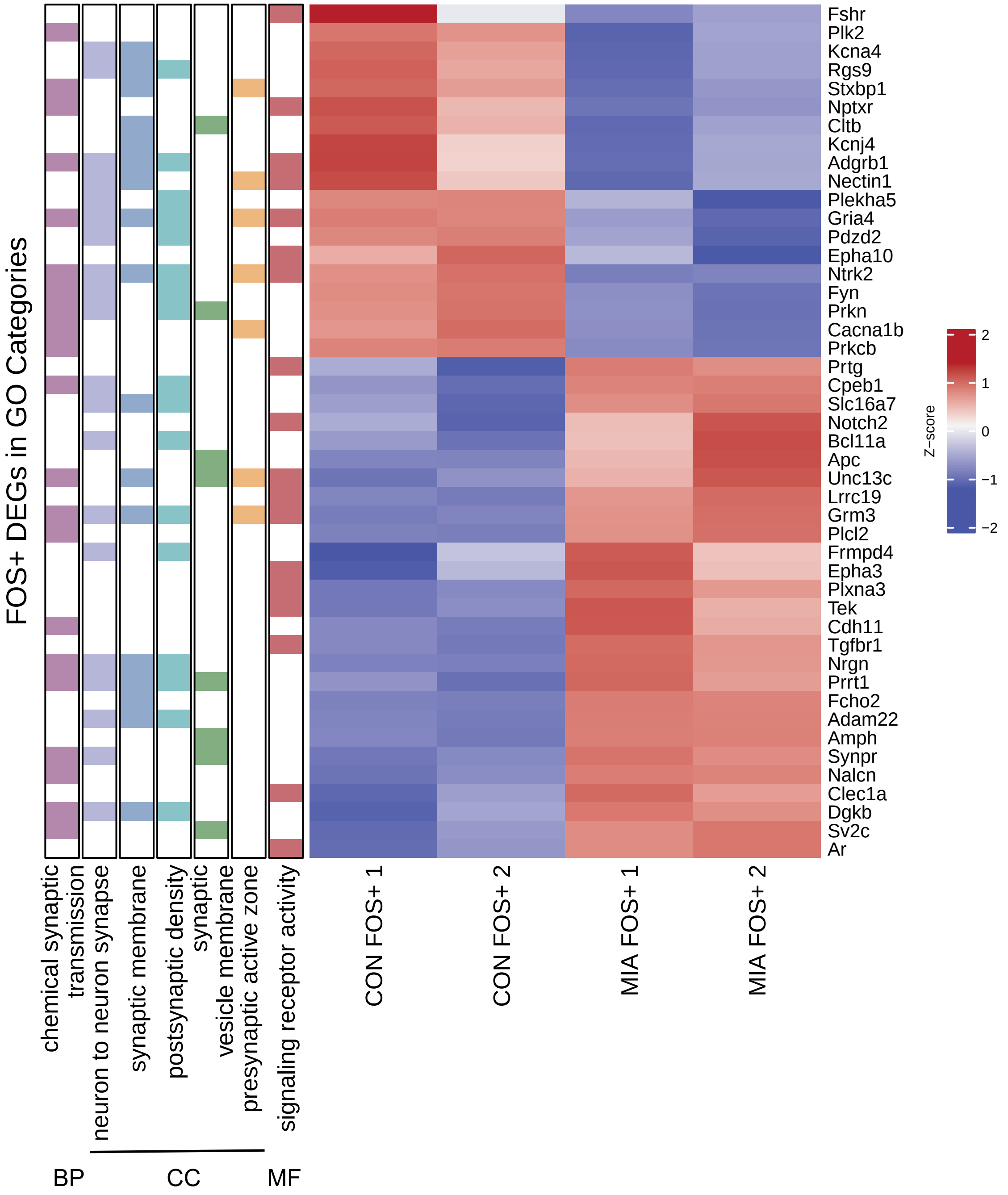


**SFigure 6.** Heatmap depicting expression of FOS+ DEGs enriched in GO categories. Genes are color-coded based on GO categories. BP, biological function; CC (see Fig. 4G), cell compartment; MF, molecular function.
