## Supplementary material for "Prenatal adversity configures a subpopulation of ventral dentate granule cells for recruitment to drive innate anxiety": Methods

### Animals

All animal experiments were carried out in accordance with the National Institutes of Health guidelines under protocols approved by the Weill Cornell Medical College Institutional Animal Care and Use Committee. All mice were group housed under a 12–hour light/dark cycle with lights on at 6 a.m. Food and water were available *ad libitum*. Experimental mice were either Swiss Webster wild type (WT) control (CON) or 5HT1A–$R^{+/+}$ (MIA) males and ≥ 12 weeks of age. 5HT1A–R deficient mice were generated as previously reported[1].

### Elevated plus maze

Mice were habituated to the behavior room for 1 hour (2 hours for snRNA-seq experiments) prior to testing. Briefly, mice were tested at 13 lux (dim overhead lighting) on a cross EPM maze with 19.5 x 2.5 inch arms. Animals were placed on the middle portion of the maze facing an open arm and allowed to freely explore for 10 min. Time spent in, number of entries into, and distance traveled in the open and closed arms were measured by the video tracking system EthoVision 15 (Noldus Information Technology). Differences between groups were assessed using unpaired t-tests and adjusted for multiple testing corrections using the Holm-Sidak method in GraphPad Prism.

### Slice electrophysiology

Slice electrophysiology experiments were performed as previously described[2]. Mice were rapidly decapitated under isoflurane anesthesia before brain collection. Brains were blocked on the coronal plane and acute 300 µm coronal slices prepared and incubated in carbogenated solutions with a pH of 7.35 and osmolarity of 305 as previously described[3]. Sections including the ventral dentate gyrus were sliced in N-methyl-D-glucamine (NMDG) artificial cerebrospinal fluid (NMDG-aCSF) at room temperature (RT) containing (in mM): 92 NMDG, 2.5 KCl, 1.25 NaH2PO4, 30 NaHCO3, 20 HEPES, 25 glucose, 2 thiourea, 5 Na-ascorbate, 3 Na-pyruvate, 0.5 CaCl2·2H2O, and 10 MgSO4·7H2O. Slices were transferred to NMDG-aCSF at 32°C for 12-14 minutes, and then incubated for at least 1 hour at RT in HEPES-aCSF containing (in mM): 92 NaCl, 2.5 KCl, 1.25 NaH2PO4, 30 NaHCO3, 20 HEPES, 25 glucose, 2 thiourea, 5 Na-ascorbate, 3 Na-pyruvate, 2 CaCl2·2H2O, and 2 MgSO4·7H2O. Slices were placed in the recording chamber and perfused at a rate of 2 mL/minute with 30°C normal aCSF containing (in mM): 124 NaCl, 2.5 KCl, 1.25 $NaH_2PO_4$, 24 $NaHCO_3$, 12.5 glucose, 5 HEPES, 2 $CaCl_2 \cdot 2H_2O$, and 2 $MgSO_4 \cdot 7H_2O$, for at least 20 minutes prior to electrophysiological recordings. Miniature postsynaptic currents were measured in a voltage clamp configuration using a cesium-gluconate-based intracellular recording solution containing (in mM): 117 D-gluconic acid, 20 HEPES, 0.4 EGTA, 5 TEA, 2 $MgCl_2$-$6H_2O$, 4 Na-ATP and 0.4 Na-GTP (pH 7.3 and 290 mOsm), and a holding potential of -70 mV. Miniature postsynaptic currents were measured in the presence of tetrodotoxin (TTX, 1 μM) in the aCSF bath. Signals were acquired using a Multiclamp 700B amplifier (Molecular Devices), digitized, and analyzed via pClamp 10.4 or 11 software (Molecular Devices). Excitability experiments were performed in current-clamp configuration using a potassium gluconate-based intracellular recording solution containing (in mM): 135 KGluc, 5 NaCl, 2 $MgCl_2$−$6H_2O$, 10 HEPES, 0.6 EGTA, 4 Na-ATP and 0.4 NaGTP (pH 7.3 and 290 mOsm) at -70 mV and resting membrane potential. Input resistance and access resistance were continuously monitored throughout experiments, and cells in which properties changed by more than 20% were not included in data analysis.

**Single nucleus isolation**

Following cervical dislocation, brains were extracted and cut coronally into 1mm sections on a brain block. The ventral hippocampus was dissected from the appropriate sections and collected into D-PBS (without $Ca^{2+}$ or $Mg^{2+}$) with 1X protease inhibitor on ice. Before nuclei isolation, the tissue was spun down and the buffer was removed. Nuclei were isolated using the Minute™ Single Nucleus Isolation Kit for Neuronal Tissues/Cells (Invent Biotechnologies, Inc.) with slight modifications to manufacturer protocol. After adding 200 ul of Buffer A, tissue was homogenized with a pestle using 30-50 twists. An additional 400 ul of Buffer A was added, and tissue was homogenized with another 10 twists and then incubated at -20°C for 10 minutes. Homogenate was first filtered through a 70 uM filter, and then transferred to a filter column provided in the kit. The homogenate was spun at 800g for 5 minutes at 4°C. The pellet was resuspended with 500 ul of 1% BSA in DPBS with 0.2U/ul of RNase inhibitor and spun again at 600 g for 5 minutes at 4C. Nuclei were resuspended in the same buffer to a concentration of 1000-5000 nuclei/ul.

**Fluorescence activated nuclei sorting**

Nuclei were stained for sorting as previously described[4]. Briefly, first, nuclei were incubated with Fc Block ((101302, Biolegend, 1:1000) for 10 minutes to reduce non-specific antibody binding on ice. Nuclei were then stained with anti-PROX1 Alexa Flour 647 (NBP-1-30045AF647, NovusBio, 1:5000) and/or anti-c-FOS Alexa Flour 488 (NBP2-50037AF488, NovusBio, 1:5000), and incubated on ice for 30 minutes. Nuclei were washed with an equal volume of buffer (1% BSA in D-PBS with 0.2 U/ul of RNase inhibitor) and spun at 600 g for 5 minutes at 4°C. Nuclei resuspended to a concentration of 1000-5000 nuclei/ul were sorted on a BD Influx sorter.

**Reduced representative bisulfite sequencing (RRBS) and analysis**

Brains were extracted after cervical dislocation, frozen on dry ice and sectioned (200 μM) on the cryostat. DNA was isolated from microdissected ventral dentate gyrus tissue using the QIAamp DNA Micro Kit (Qiagen) according to manufacturer instructions. RRBS libraries were prepared by the Epigenomics Core at Weill Cornell Medicine. Libraries were sequenced on a NovaSeq6000 in a 100 bp paired end mode with a target of 100 million reads per sample (n = 4 animals per group).

Alignment to the mm10 reference genome and methylation calling was done using Bismark v0.22.3[5]. Differential methylation analysis was performed with methylKit v1.26 in R 4.3.1[6]. Briefly, CGs with at least 10 coverage reads but not more than the 99.9 percentile of coverage reads were kept. Samples were merged, and only CGs covered by all samples were tested for differential methylation using default settings. CGs were considered differentially methylated with 1) a significance threshold of $q < 0.01$ and 2) a 10% difference in methylation between the groups of interest. Differentially methylated regions (DMRs) were defined as regions containing at least two differentially methylated CpGs with ≤ 1 kb distance between the sites. DMRs were classified as hypomethylated, hypermethylated or mixed based on whether all differentially methylated CpGs in the DMR were hypomethylated, hypermethylated or a combination of both. Genes were assigned to DMRs using rGREAT[7].

Background block regions were determined by clustering all tested CG sites with the same criteria used to cluster DMRs.

**DMR annotation**

Mm10 genomic coordinates for exons and introns were downloaded from the University of California Santa Cruz (UCSC) genome browser. Promoters were defined as regions ± 500 bp from the transcription start site (TSS). Intergenic regions were found through exclusion of genic regions from chromosome coordinates. DMR enrichments in chromatin states[8] were calculated using a Fisher's exact test comparing against background block regions. SEA from MEME Suite tools was used for motif enrichment analysis[9,10].

**Single nucleus RNA sequencing (snRNA–seq) and analysis**

Final single nuclei suspensions (n = 2 per sample group each pooled from 6 animals) were submitted to the Weill Cornell Epigenomic Core for library preparation and sequencing. Sample libraries were prepared using the Chromium Single Cell Reagent V3 Kit from 10X Genomics according to manufacturer instructions. Libraries were sequenced on an Illumina NovaSeq to a depth of approximately 25,000–30,000 target reads per nucleus.

Raw sequencing data were processed using the CellRanger v6.0.0 pipeline and aligned to the mm10 transcriptome reference with introns included. Gene–expression matrices of samples were loaded into R, assessed for quality, and analyzed using Seurat[11]. Nuclei with fewer than 500 or greater than 6000 genes, greater than 0.1% mitochondrial reads gene reads, and fewer than 1000 UMIs were excluded from analysis. Genes expressed in fewer than 10 nuclei were also filtered out. After normalization with Sctransform v2, samples were integrated using 3,000 variable features. Nuclei were clustered using 40 PCs with a resolution of 0.5. 2 low quality clusters were removed after which nuclei were re-clustered using 40 PCs and a resolution of 0.6. One FOS+ cluster was subclustered using a resolution of 0.1. Cluster markers were found with the FindAllMarkers function (min.pct = 0.25, logfc.threshold = 0.25) using the default Wilcoxon rank sum test with Bonferroni correction, and cell types were annotated using known cell markers.

For differential expression testing, DGC clusters were subset and counts were aggregated based on the following sample identities: CON FOS-, CON FOS+, MIA FOS-, MIA FOS+. Pseudobulk samples were analyzed for differential expression using DESeq2[12] using default parameters. Genes with at least 50 aggregated counts in at least two samples were tested. Differential expression gene lists (p-adjusted < 0.1) were filtered to keep only genes expressed in at least 5% of one sample group tested. Overrepresentation analysis for enrichment in gene ontology categories was done using ClusterProfiler[13] with all tested genes set as the background universe. Significant categories (p-adjusted < 0.05) were then filtered for redundancy based on p-value and genes represented in each category. Differentially expressed genes that are also neuropsychiatric disease risk genes were identified with psygenet2r[14].

**Intracranial fiber implantation into the vDG for fiber photometry**.

Male animals were anesthetized with 2% isoflurane and placed onto a stereotaxic frame to perform craniotomy. First, we delivered 275 nL viral vector AAV1.Syn.GCaMP6s at a rate of 40 nL per minute by a 1 µL Hamilton 7000 series syringe with a 32 gauge blunt needle attached to a microsyringe pump and controller at the following coordinates: -3.65 mm anterior-posterior, -2.7 mm medial-lateral, and -3.0 mm dorsal-ventral. The needle remained at the injection site for 7 minutes post injection and was withdrawn slowly. A 400 µm diameter optical fiber was then implanted above the vDG using the following coordinates: -3.65 mm anterior-posterior, -2.7 mm medial-lateral, and -2.8 mm dorsal-ventral. Optical fibers were secured with Metabond.

**Fiber photometry and data analysis**

Optical patch cables were photobleached through the fiber photometry workstation (Tucker-Davis Technologies, Inc., Alachua, Florida) for at least 12 h prior to being tethered to the mouse's head stage prior to the trial. During each 10-min EPM trial, raw GCaMP and isosbestic control signals were collected in the 465 nm and 405 nm channels, respectively, at a sampling rate of 1017.25 Hz. The acquired signals were processed via a custom Python script in Jupyter Notebook open-source software. The signals were first denoised by removing electrical artifacts using a median filter and by reducing high-frequency noise with a lowpass filter, followed by downsampling by factor of 100. Resulting preprocessed data were corrected for motion artifacts and photobleaching by finding a linear fit of the GCaMP and isosbestic control signal, scaling the isosbestic control by the resulting coefficients of the linear fit, and subtracting the scaled control signal from the GCaMP signal. Final signal values were calculated by computing a z-score based on the data from the whole trial. Transient events within the GCaMP signal were detected using the Mini Analysis program Version 6. 0. 3 (Synaptosoft, USA). To identify parts of the final GCaMP signal that correspond to the animal's location in different sections of the EPM arena, we tracked animals' position during the trials using Ethovision 15 software (Noldus, Wageningen, Netherlands). Lastly, correct localization and extent of viral expression were verified in each mouse using fluorescent microscopy.
